# Infection and vaccine-induced neutralizing antibody responses to the SARS-CoV-2 B.1.617.1 variant

**DOI:** 10.1101/2021.05.09.443299

**Authors:** Venkata-Viswanadh Edara, Lilin Lai, Malaya K. Sahoo, Katharine Floyd, Mamdouh Sibai, Daniel Solis, Maria W. Flowers, Laila Hussaini, Caroline Rose Ciric, Sarah Bechnack, Kathy Stephens, Elham Bayat Mokhtari, Prakriti Mudvari, Adrian Creanga, Amarendra Pegu, Alexandrine Derrien-Colemyn, Amy R. Henry, Matthew Gagne, Barney S. Graham, Jens Wrammert, Daniel C. Douek, Eli Boritz, Benjamin A. Pinsky, Mehul S. Suthar

## Abstract

SARS-CoV-2 has caused a devastating global pandemic. The recent emergence of SARS-CoV-2 variants that are less sensitive to neutralization by convalescent sera or vaccine-induced neutralizing antibody responses has raised concerns. A second wave of SARS-CoV-2 infections in India is leading to the expansion of SARS-CoV-2 variants. The B.1.617.1 variant has rapidly spread throughout India and to several countries throughout the world. In this study, using a live virus assay, we describe the neutralizing antibody response to the B.1.617.1 variant in serum from infected and vaccinated individuals. We found that the B.1.617.1 variant is 6.8-fold more resistant to neutralization by sera from COVID-19 convalescent and Moderna and Pfizer vaccinated individuals. Despite this, a majority of the sera from convalescent individuals and all sera from vaccinated individuals were still able to neutralize the B.1.617.1 variant. This suggests that protective immunity by the mRNA vaccines tested here are likely retained against the B.1.617.1 variant. As the B.1.617.1 variant continues to evolve, it will be important to monitor how additional mutations within the spike impact antibody resistance, viral transmission and vaccine efficacy.

---

SARS-CoV-2 has caused a devastating global pandemic. The recent emergence of SARS-CoV-2 variants that are less sensitive to neutralization by convalescent sera or vaccine-induced neutralizing antibody responses has raised concerns^1,2^. A second wave of SARS-CoV-2 infections in India is leading to the expansion of SARS-CoV-2 variants. These variants contain mutations arising within the spike protein that are known to increase resistance to antibody neutralization^3^. The B.1.617.1 variant was first identified in India and has rapidly spread throughout India and to several countries throughout the world. In this study, using a live virus assay, we describe the neutralizing antibody response to the B.1.617.1 variant in serum from infected and vaccinated individuals.

The B.1.617.1 was isolated from a residual mid turbinate swab from a patient in Stanford, CA in March 2021 (hCoV-19/USA/CA-Stanford-15_S02/2021). Relative to the WA1/2020 virus (nCoV/USA_WA1/2020), the B.1.617.1 variant contains several mutations within the spike protein, including within the N-terminal antigenic supersite (G142D and E154K)^4^, the receptor binding domain (L452R and E484Q) and within the polybasic furin cleavage site at the S1/S2 boundary (P681R). Here, we used a Live virus Focus Reduction Neutralization Test (FRNT)^5^ to compare the neutralizing antibody response in serum from 24 convalescent COVID-19 individuals (31-91 days after symptom onset)^1^, 15 mRNA-1273 vaccinated individuals (35-51 days post-2nd dose), and 10 BNT162b2 vaccinated individuals (7-27 days post-2nd dose).

Across samples from infected and vaccinated individuals, all individuals showed reduced neutralization titers against the B.1.617.1 variant. In the convalescent sera samples, the FRNT_50_ geometric mean titers (GMT) were 514 for WA1/2020 (95% CI, 358 to 740) and 79 for B.1.617.1 (95% CI, 49 to 128), and 5 samples were undetectable against the B.1.617.1 variant. Among the mRNA-1273 vaccinated sera samples, the GMTs were 1332 for WA1/2020 (95% CI, 905 to 1958) and 190 for B.1.617.1 (95% CI, 131 to 274). In the BNT162b2 vaccinated sera samples, the GMTs were 1176 for WA1/2020 (95% CI, 759 to 1824) and 164 for B.1.617.1 (95% CI, 104 to 258). Among the three sample groups the FRNT_50_ GMTs for B.1.617.1 were statistically significantly lower than the WA1/2020 strain.

Our results show that the B.1.617.1 variant is 6.8-fold less susceptible to neutralization by sera from infection and vaccinated individuals. Despite this, a majority of the sera from convalescent individuals (79%; 19/24 samples) and all sera from vaccinated individuals were still able to neutralize the B.1.617.1 variant. This suggests that protective immunity by the mRNA vaccines tested here are likely retained against the B.1.617.1 variant. As the B.1.617.1 variant continues to evolve, it will be important to monitor how additional mutations within the spike impact antibody resistance, viral transmission and vaccine efficacy.

**Figure 1:**
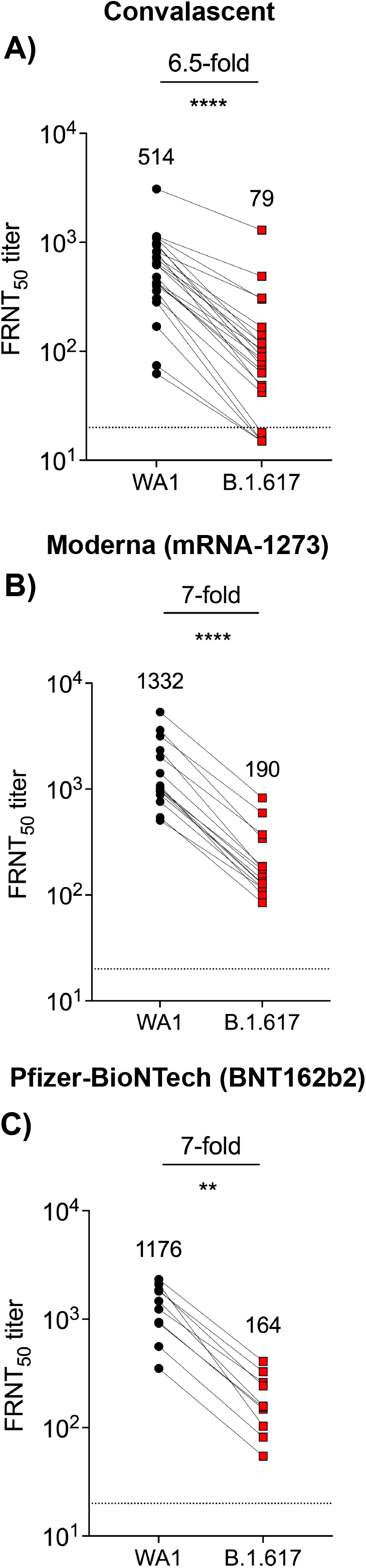
Neutralizing antibody responses between WA1/2020 and B.1.617.1 viruses after infection and vaccination. Data from the following cohorts are shown from natural infection: 24 convalescent COVID-19 individuals (31-91 days after symptom onset, panel A), Moderna (mRNA-1273) vaccinated: 15 individuals (35-51 days post-2nd dose, panel B), and Pfizer-BioNTech (BNT162b2) vaccinated: 10 individuals (7-27 days post-2nd dose, panel C). In panels A-C, the FRNT_50_ GMTs for WA1/2020 and B.1.617.1 are shown. The connecting lines between WA1/2020 and B.1.617.1 represents matched serum samples. The horizontal dashed lines along the X-axis indicate the limit of detection (FRNT50 GMT= 20). Normality of the data was determined using Shapiro Wilk normality test. Non-parametric pairwise analysis for neutralization titers were performed by Wilcoxon matched-pairs signed rank test. **p<0.01; ****p<0.0001

## Acknowledgments

This work was supported in part by grants (NIH P51 OD011132, 3U19AI057266-17S1 to Emory University) from the National Institute of Allergy and Infectious Diseases (NIAID), National Institutes of Health (NIH), by intramural funding from the National Institute of Allergy and Infectious Diseases, by The Oliver S. and Jennie R. Donaldson Charitable Trust, Emory Executive Vice President for Health Affairs Synergy Fund award, the Pediatric Research Alliance Center for Childhood Infections and Vaccines and Children’s Healthcare of Atlanta, the Emory-UGA Center of Excellence for Influenza Research and Surveillance, COVID-Catalyst-I^3^ Funds from the Woodruff Health Sciences Center and Emory School of Medicine, Woodruff Health Sciences Center 2020 COVID-19 CURE Award. Funders played no role in the design and conduct of the study; collection, management, analysis, and interpretation of the data; preparation, review, or approval of the manuscript; and decision to submit the manuscript for publication.

