## Supplementary appendix for "Infection and vaccine-induced neutralizing antibody responses to the SARS-CoV-2 B.1.617.1 variant"

### Supplemental Methods

Viruses and cells**.** VeroE6-TMPRSS2 cells were generated by transfecting VERO E6 cells (ATCC CRL-1586) with pCAGGS plasmid in which chicken actin gene promoter drives the expression of an open reading frame comprising Puromycin N-acetyl transferase, GSG linker, 2A self-cleaving peptide of thosea asigna virus (T2A), human transmembrane serine protease 2 (TMPRSS2). Two days post-transfection, cells were trypsinzed and transferred to a 100 mm dish containing complete DMEM medium (1x DMEM, Thermo Fisher, # 11965118, 10% FBS, 1x penicillin/streptomycin) supplemented with puromycin (Thermo Fisher, #A1113803) at a final concentration of 10 µg/ml. Approximately ten days later, individual colonies of cells were isolated using cloning cylinders (Sigma) and expanded in medium containing puromycin. Clonal cell lines were screened for expression of TMPRSS2 by flow cytometry. VeroE6-TMPRSS2 cells were cultured in complete DMEM in the presence of Gibco Puromycin 10mg/mL (# A11138-03). nCoV/USA_WA1/2020 (WA/1), closely resembling the original Wuhan strain and resembles the spike used in the mRNA-1273 and Pfizer BioNTech vaccine, was propagated from an infectious SARS-CoV-2 clone as previously described^1^. icSARS-CoV-2 was passaged once to generate a working stock. hCoV-19/USA/CA-Stanford-15_S02/2021 (herein referred to as the B.1.617 variant) was derived from a mid-turbinate nasal swab collected from a Stanford Healthcare patient in March 2021, as part of a study approved by the institutional review board at Stanford University (Protocol 57519). As described previously^2,3^, viral genome enrichment was conducted using laboratory-developed, multiplex RT-PCR reactions that generate multiple overlapping amplicons ~1200 base-pairs in length. Fragment libraries were prepared using NEBNext DNA Library Prep reagents for Illumina (New England BioLabs, Ipswich, MA), and were sequenced on an Illumina MiSeq using single-end 150 cycle sequencing using MiSeq Reagent kit V3. The genome was assembled via a custom assembly and bioinformatics pipeline using NCBI NC_045512.2 as reference. 724x mean whole genome coverage was obtained for this sample; 100x coverage was obtained over 93.4% of the genome and 99.9% of spike. The consensus SARS-CoV-2 genome is available under GISAID accession number EPI_ISL_1675223. The B.1.617 variant was plaque purified on VeroE6-TMPRSS2 cells and propagated once in a 12-well plate of confluent VeroE6-TMPRSS2 cells followed by expansion of the working stock in a T175 flask of confluent VeroE6-TMPRSS2 cells. The working stock was aliquoted to generate a working stock and deep sequenced (**Supplementary Table 1**). Illumina-ready libraries were generated using NEBNext Ultra II RNA Prep reagents (New England BioLabs) as previously described^4^. Briefly, we fragmented RNA, followed by double-stranded cDNA synthesis, end repair, and adapter ligation. The ligated DNA was then barcoded and amplified by a limited cycle PCR and the barcoded Illumina libraries were sequenced by using paired-end 150-base protocol on a NextSeq 2000 (Illumina). Demultiplexed sequence reads were analyzed in the CLC Genomics Workbench v.21.0.3 by (i) trimming for quality, length, and adaptor sequence, (ii) mapping to the Wuhan-Hu-1 SARS-CoV-2 reference (GenBank accession number: NC_045512), (iii) improving the mapping by local realignment in areas containing insertions and deletions (indels), and (iv) generating both a sample consensus sequence and a list of variants. Default settings were used for all tools.

Samples. For samples from Emory University, collection and processing were performed under the University Institutional Review Board protocols #00001080 and #00022371. Adults ≥18 years were enrolled who met eligibility criteria for SARS-CoV-2 infection and provided informed consent. Convalescent samples were a convenience sample of individuals that had recovered from mild or moderate COVID-19. Patients had PCR or rapid antigen-test confirmed COVID-19 between the months of March-August 2020 and enrolled with samples collected from May-October 2020. These individuals were recruited using multiple methods, including advertisements on the university campus, primary care clinics and at COVID-19 testing sites. Interested participants contacted the clinical research site and underwent a phone screening to assess if they met eligibility criteria. In addition, primary care clinic patients who were being managed for COVID-19 were contacted to see if they are interested in participating in this study. At the time of study enrollment, some of the participants had residual symptoms but others had recovered with no residual symptoms. No data was collected on the number of patients that were pre-screened or declined participation. The Pfizer serum samples were deposited by Florian Krammer, Ph.D. and Viviana Simon, Ph.D., Icahn School of Medicine at Mount Sinai, New York, New York, USA and was obtained through BEI Resources, NIAID, NIH: SARS-CoV-2 Vaccine Human Serum, NR-55279. Plasma was collected from volunteers, previously vaccinated with the Moderna vaccine, at the Emory Hope Clinic and Emory Children’s Center under IRB approvals 00045821 and 00022371, respectively. The study enrolled and obtained informed consent from adult participants who had received the complete schedule of the Moderna vaccine. The description of the serum samples from convalescent individuals is shown in **Supplementary Table 2** and from Moderna (mRNA-1273) vaccinated individuals in **Supplementary Table 3** and from Pfizer-BioNTech (BNT162b2) vaccinated individuals in **Supplementary Table 4**.

Focus Reduction Neutralization Test**.** FRNT assays were performed as previously described^5^. Briefly, samples were diluted at 3-fold in 8 serial dilutions using DMEM (VWR, #45000-304) in duplicates with an initial dilution of 1:10 in a total volume of 60 μl. Serially diluted samples were incubated with an equal volume of WA1/2020 or B.1.617 (100-200 foci per well based on the target cell) at 37^o^ C for 1 hour in a round-bottomed 96-well culture plate. The antibody-virus mixture was then added to VeroE6-TMPRSS2 cells and incubated at 37^o^ C for 1 hour. Post-incubation, the antibody-virus mixture was removed and 100 µl of pre-warmed 0.85% methylcellulose (Sigma-Aldrich, #M0512-250G) overlay was added to each well. Plates were incubated at 37^o^ C for 16 hours. After 16 hours, methylcellulose overlay was removed, and cells were washed three times with PBS. Cells were then fixed with 2% paraformaldehyde in PBS for 30 minutes. Following fixation, plates were washed twice with PBS and 100 µl of permeabilization buffer, was added to the fixed cells for 20 minutes. Cells were incubated with an anti-SARS-CoV spike primary antibody directly conjugated with alexaflour-488 (CR3022-AF488) for up to 4 hours at room temperature. Cells were washed three times in PBS and foci were visualized and imaged on an ELISPOT reader (CTL).

Quantification and Statistical Analysis. Antibody neutralization was quantified by counting the number of foci for each sample using the Viridot program^6^. The neutralization titers were calculated as follows: 1 - (ratio of the mean number of foci in the presence of sera and foci at the highest dilution of respective sera sample). Each specimen was tested in duplicate. The FRNT-50 titers were interpolated using a 4-parameter nonlinear regression in GraphPad Prism 8.4.3. Samples that do not neutralize at the limit of detection at 50% are plotted at 15 and was used for geometric mean calculations.

**Supplemental Figure 1.**

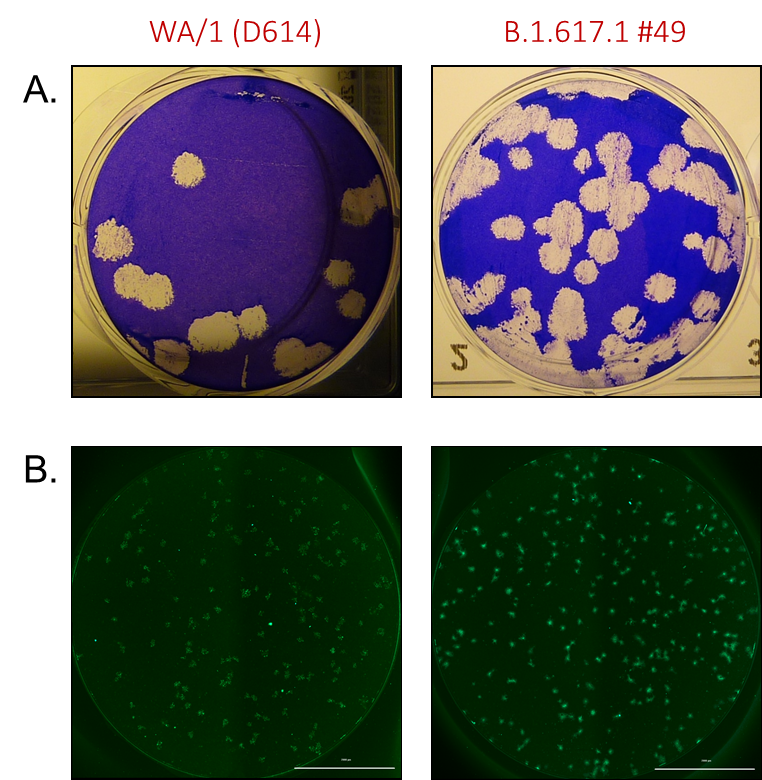

Supplemental Figure 1. Plaque and focus-forming morphology of WA1/2020 and B.1.617**.** Monolayers of VeroE6-TMPRSS2 cells were infected with SARS-CoV-2 strains WA1/2020 and B.1.617. A. Plaque morphology- At 60 hours post-infection, monolayers were fixed with 4% paraformaldehyde for 30 minutes and stained with 0.1% crystal violet in 20% methanol for 10 minutes. were plaqued on a monolayer of VeroE6-TMPRSS2 cells. B. Foci morphology- Focus forming assay was performed as described in the methods. At 16 hours post-infection, monolayers were fixed with 2% paraformaldehyde for 30 minutes, washed with PBS, permeabilized, and stained with a AF488-conjugated CR3022 monoclonal antibody. Cells were visualized using an ELISPOT reader.

### **Supplemental Table 1**. Nucleotide variants and Amino acid mutations identified by deep sequencing results of the B.1.617 working stock

| **Type** | **Variant** | **Amino Acid Mutation** | **Gene (Region)** |
| --- | --- | --- | --- |
| SNP | 3037C>T | -- | orf1ab |
| SNP | 3457C>T | -- | orf1ab |
| SNP | 4780C>T | -- | orf1ab |
| SNP | 4965C>T | T1567I | orf1ab (nsp3) |
| SNP | 5907C>T | T1881I | orf1ab (nsp3) |
| SNP | 11201A>G | T3646A | orf1ab (nsp6) |
| SNP | 14408C>T | P4715L | orf1ab (nsp12) |
| SNP | 16362C>T | -- | orf1ab |
| SNP | 17523G>T | M5753I | orf1ab (nsp13) |
| SNP | 18511C>T | -- | orf1ab |
| SNP | 20016C>T | -- | orf1ab |
| SNP | 20396A>G | K6711R | orf1ab (nsp15) |
| SNP | 20936C>T | T6891M | orf1ab (nsp16) |
| SNP | 21895T>C | -- | spike |
| SNP | 21987G>A | G142D | spike |
| SNP | 22022G>A | E154K | spike |
| SNP | 22917T>G | L452R | spike |
| SNP | 23012G>C | E484Q | spike |
| SNP | 23403A>G | D614G | spike |
| SNP | 23604C>G | P681R | spike |
| SNP | 24775A>T | Q1071H | spike |
| SNP | 24863C>G | H1101D | spike |
| SNP | 25276C>T | -- | spike |
| SNP | 25469C>T | S26L | orf3a |
| SNP | 26256C>A | F4L | E |
| SNP | 27299T>C | I33T | orf6 |
| SNP | 27638T>C | V82A | orf7a |
| SNP | 28881G>T | G202W | N |
| SNP | 29402G>T | -- | N |

### **Supplemental Table 2**. FRNT50 results from convalescent COVID-19 samples

| **Sample#** | **Date samples collected** | **Days After Symptom Onset** | **WA1/2020 FRNT50** | **B.1.617 FRNT50** |
| --- | --- | --- | --- | --- |
| 1 | 4/17/20 | 34 | 359 | <20 |
| 2 | 4/22/20 | 44 | 479 | 49 |
| 3 | 4/22/20 | 31 | 423 | 105 |
| 4 | 4/22/20 | 37 | 74 | <20 |
| 5 | 4/23/20 | 39 | 305 | 42 |
| 6 | 4/23/20 | 40 | 1129 | 487 |
| 7 | 4/27/20 | 30 | 283 | <20 |
| 8 | 4/28/20 | 46 | 716 | 100 |
| 9 | 4/28/20 | 42 | 1101 | 300 |
| 10 | 4/29/20 | 50 | 1088 | 116 |
| 11 | 4/30/20 | 48 | 975 | 72 |
| 12 | 5/6/20 | 47 | 820 | 47 |
| 13 | 5/8/20 | 56 | 622 | 119 |
| 14 | 5/13/20 | 61 | 646 | 143 |
| 15 | 5/15/20 | 60 | 169 | <20 |
| 16 | 5/21/20 | 57 | 400 | 87 |
| 17 | 7/29/20 | 31 | 62 | <20 |
| 18 | 8/21/20 | 52 | 626 | 80 |
| 19 | 8/14/20 | 83 | 3085 | 1296 |
| 20 | 8/21/20 | 50 | 944 | 89 |
| 21 | 8/21/20 | 76 | 425 | 67 |
| 22 | 8/26/20 | 58 | 423 | 63 |
| 23 | 8/26/20 | 42 | 819 | 308 |
| 24 | 10/9/20 | 91 | 734 | 167 |

### Supplemental Table 3. FRNT50 results from Moderna (mRNA-1273) vaccinated samples

| **Sample #** | **Date samples collected** | **Days After Second Dose** | **WA1/2020 FRNT50** | **B.1.617 FRNT50** |
| --- | --- | --- | --- | --- |
| 1 | 3/25/21 | 48 | 956 | 173 |
| 2 | 3/25/21 | 49 | 540 | 85 |
| 3 | 3/25/21 | 47 | 957 | 126 |
| 4 | 3/25/21 | 50 | 884 | 140 |
| 5 | 3/25/21 | 47 | 505 | 117 |
| 6 | 3/25/21 | 47 | 1074 | 100 |
| 7 | 3/25/21 | 35 | 3593 | 340 |
| 8 | 3/26/21 | 50 | 1014 | 159 |
| 9 | 3/26/21 | 48 | 2322 | 175 |
| 10 | 3/26/21 | 38 | 5330 | 826 |
| 11 | 3/26/21 | 50 | 759 | 118 |
| 12 | 3/26/21 | 51 | 1012 | 128 |
| 13 | 3/26/21 | 50 | 1411 | 186 |
| 14 | 3/26/21 | 48 | 2007 | 372 |
| 15 | 3/26/21 | 50 | 3159 | 595 |

### Supplemental Table 4. FRNT50 results from Pfizer -BioNTech (BNT162b2) vaccinated samples

| **Sample #** | **Days After Second Dose** | **WA1/2020 FRNT50** | **B.1.617 FRNT50** |
| --- | --- | --- | --- |
| 1 | 6 | 1237 | 261 |
| 2 | 7 | 937 | 148 |
| 3 | 27 | 560 | 82 |
| 4 | 8 | 1859 | 242 |
| 5 | 23 | 1812 | 327 |
| 6 | 22 | 351 | 55 |
| 7 | 22 | 916 | 154 |
| 8 | 20 | 2106 | 103 |
| 9 | 9 | 2331 | 408 |
| 10 | 21 | 1468 | 158 |
